# Fatigue, effort perception and central activation failure in chronic stroke survivors: a TMS and fMRI investigation

**DOI:** 10.1101/821959

**Authors:** Isobel F Turner, Nick S Ward, Annapoorna Kuppuswamy

## Abstract

Fatigue is commonly seen in stroke survivors and the most common manifestation of fatigue is the requirement of high effort for activities of daily life. In this study we set out to identify the neural correlates of perceived effort and central activation failure, a neurophysiological measure correlated with perceived effort. Methods: Twelve chronic stroke survivors participated in this study. Fatigue levels were quantified using the Fatigue Severity Scale -7, perceived effort was quantified using a 1-10 numerical rating scale while performing an isometric biceps hold task, Central Activation Failure was quantified using the modified twitch interpolation technique using Transcranial Magnetic Stimulation and functional Magnetic Resonance Imaging was used to measure blood-oxygen-level dependent signal (BOLD) from the brain while the participant performed a hand grip task. Analysis: Following standard pre-processing procedures for fMRI data using SPM software, co-variance of BOLD signal with perceived effort levels and central activation failure was evaluated. Correlation analysis was performed between measures of fatigue and effort. Results: The main findings of this study were 1) high fatigue was associated with high perceived effort 2) higher perceived effort was associated with greater increase in BOLD fMRI activity in pre-SMA and the ipsilateral inferior frontal gyrus with increasing force 3) greater Central Activation Failure was associated with higher increase in BOLD fMRI activity in bilateral caudate, contralateral superior frontal gyrus and pre-motor cortices with increasing force.

## Introduction

Fatigue is common after stroke and little is known about its pathophysiology. One of the most common manifestations of fatigue is the exaggerated notion of effort required to perform activities of daily living when compared to the pre-stroke condition. Effort in the context of physical activity is the subjective sense of muscular force and is thought to arise from neural components that drive voluntary motor output. Increased neural activity with heightened effort perception has been noted in the prefrontal cortices [1], motor and secondary motor areas [2,3]. Effort related neural activity precedes muscle activation [3] suggesting a non-dependence of effort perception on sensory afferent information arising from the activated muscle. Converging evidence from reports of effort perception from deafferented patients [4], paralysed patients [5] and healthy fatigued subjects [2] add further support to central origins of effort perception.

In stroke survivors, despite several studies alluding to fatigue being associated with activities being more effortful [6,7] there has been little systematic investigation into central neural correlates of effort perception in stroke survivors. A recent study suggests that stroke survivors perceive activities that require low levels of muscular force to be more effortful than healthy controls [8] and we recently showed that stroke survivors who perceived production of low levels of muscle force to be more effortful had higher levels of central activation failure, a measure of excitability of inputs that drive motor output [9]. In this study we hypothesised that 1) those with high fatigue exhibit high perceived effort 2) perceived effort and central activation failure co-varies with neural activity in secondary motor areas and basal ganglia.

## Methods

### Study design

Cross-sectional observational study. The study was approved by the Riverside Research Ethics Committee (12/LO/1474). Stroke survivors were recruited consecutively via the Thames Stroke Research Network from the University College NHS Trust Hospital and from community stroke groups.

### Patient characteristics

Twelve stroke survivors (55.22 years ± 10.9, 2 females, 36.22 months post stroke ± 55.6) with a first-time ischaemic or haemorrhagic lesion participated in the study. Patients were screened for compatibility with TMS procedures. Additional exclusion criteria included, centrally acting medication, high score on Hospital Anxiety and Depression Scale (>11) and poor function. Functional screening included upper limb functional tests and cognitive tests. Poor upper limb function was defined as having less than 60% of the unaffected limb score in more than one of the following measures: a) Nine Hole Peg Test, b) Action Research Arm Test (ARAT), and c) Grip strength. Poor cognitive function was defined as a score of more than 5 on Sustained Attention Index (SAI) and Symbol Digit Modalities Test

### Experimental procedures

All subjects participated in a two part single session study. In the first part subjects were scanned in an MRI scanner and in the second part they underwent brain stimulation using transcranial magnetic stimulation.

### Self-reported measures

Fatigue score: Fatigue Severity Scale – 7, a validated fatigue scale [10] was used to quantify fatigue. Perception of Effort -A Numerical Rating Scale of 1-10 with 10 being maximal effort and 1 being minimal effort was used to measure participants’ perception of effort while performing submaximal isometric contractions (25% MVC) of the elbow flexor muscles in a custom built force device.

### Functional Magnetic Resonance Imaging protocol

A 3T Siemens ALLEGRA scanner was used to procure both T1 weighted anatomical and T2 weighted functional images. The scanner protocol has been described in detail previously [11]. During scanning, all participants performed a series of visually cued dynamic isometric hand grips with their affected hand using an MRI compatible manipulandum as described previously [12,13]. The scan session comprised 50 hand grips (10 in each condition 15%, 25%, 35%, 45% and 55% MVC). The force was monitored throughout the scan session and feedback provided to the participants.

### Transcranial Magnetic Stimulation procedure

Voluntary Activation (VA) using modified twitch interpolation technique: The experimental set up used for obtaining measures of VA was replicated from a previous study [14]. Force and EMG measures were obtained from the affected elbow flexor group of muscles using the software Spike version 2.16. Superimposed twitches were obtained at 25%, 50%, 75% and 100% MVC using a 90mm circular coil placed over the vertex. 6 repeats at each of the conditions were performed in a random order.

## Statistical analysis

### First level analysis

#### fMRI data

All data was analysed using Statistical Parametric Mapping (SPM8, Wellcome department of Imaging Neuroscience, UK) implemented in Matlab R2013a (The Mathworks Inc., USA). The pre-processing procedure was performed as described previously [11]. For single subject analysis, two covariates, grip (Bg) and force (Bf) were defined as previously described [11]. The statistical parametric maps of the t statistic resulting from linear contrasts of each covariate was generated and stored as separate images for each subject.

#### Fatigue score

The average score of the seven statement scores were calculated and taken as the FSS-7 score. Perception of Effort -The NRS scores of 1-10 were normalised by subtracting the given score from the expected score. Therefore, a negative value represents high effort perception and positive value represents low effort perception.

Central Activation Failure – Superimposed twitch size was calculated as the difference between background force (3ms before delivery of TMS pulse) and the peak force obtained after the TMS pulse. The twitch sizes were calculated at 50%, 75% and 100% MVC. 1-SIT/BF*100 where BF-background force was used to calculate VABG. A 100% voluntary activation would imply no deficit in central activation and therefore 0 Central Activation Failure. The CAF measure was calculated by subtracting VABG from 100.

### Second level analysis

The data for the second stage of analysis comprised the pooled parameter estimates for each covariate across all subjects. Contrast images containing these data for each subject were entered into one sample t-test for each covariate of interest. After characterising the average group effects, we examined for the influence of fatigue (FSS-7), central activation failure measure (CAF) and perceived effort measure (PE) on the parameter estimates (Bg or Bf). Simple linear regression analysis was performed using SPM8, in which two orthogonal covariates were a) contrast images for each subject for the effect of interest (Bg or Bf) and b) a single value representing either CAF or PE for each subject (mean corrected and normalised across the group). SPM{t}s representing brain regions in which there is a linear relationship between either parameters and CAF or PE was generated. For each significant voxel the R^2^ for the plot of parameter estimate against CAF or PE was also calculated to illustrate the relationship.

ROI analysis: Five a priori regions of interests were defined, bilateral striatum, bilateral PmD and SMA. The ROIs consisted of 10mm diameter spheres centred on the following coordinates derived from our previous work [12,13] and with careful identification of anatomical structures with the aid of the atlas of Duvernoy (1991).

Contralateral PmD (−24, -12, 66) ipsilateral PmD (24, -12, 66), contralateral striatum (−12, -13, 19) ipsilateral striatum (12, -13, 19) and pre-SMA (2, 20, 50).

All SPM{t}s were transformed to the unit normal Z-distribution to create a statistical parametric map (SPM {Z}). All statistical t-tests carried out in SPM was one-tailed.

Pearson Product Moment correlation analysis was performed for the following sets of data: FSS-7, PE and CAF scores.

## Results

### Behavioural results

The average fatigue score of the group was 3.35 ±2.2 (Mean ±SD). The range of fatigue levels was 1-7. The lowest possible score being 0 and highest being 7. Therefore stroke survivors with both very low and very high fatigue were included. Of the 12 participants 4 overestimated the effort required to produce a 25% MVC (3 on a scale of 0-10). Of the remaining 8, seven underestimated the effort required (2 on a scale of 0-10) and one participant grossly underestimated the effort (1 on a scale of 0-10). The Central Activation Failure was low as a group (5.01 % ± 3.9 %). This suggests that on average participants were able to voluntarily activate 95% of their maximal motor output.

### Correlation analysis of self-reported measures

A significant correlation between fatigue scores and PE scores was observed. Higher the fatigue scores, higher was the perceived effort i.e. those with high fatigue exhibited negative PE error score and those with low fatigue had positive PE error score (Figure 1), CC = -0.679, p = 0.0151.

**Figure 1:**
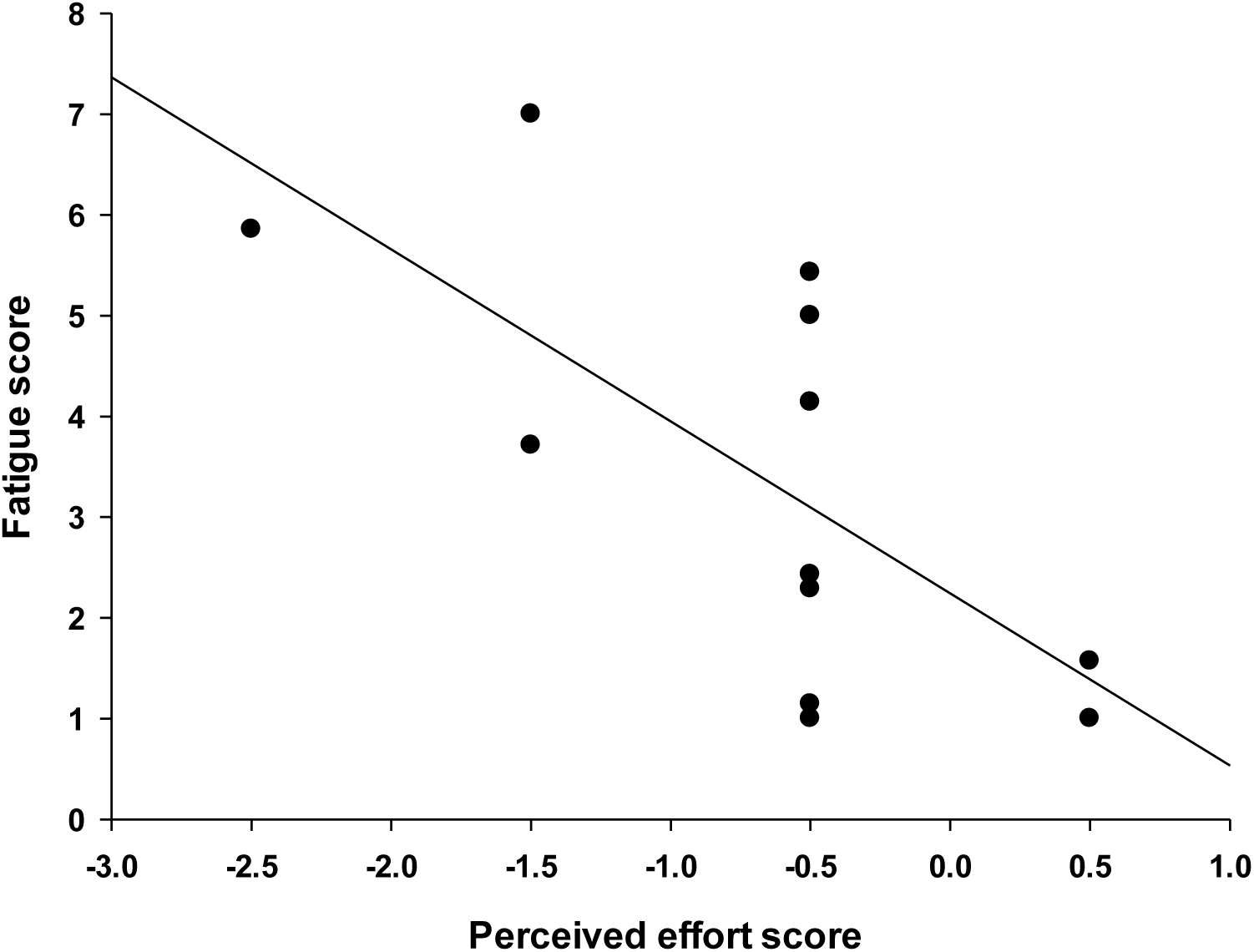
The relationship between fatigue score and perceived effort is plotted in this figure, where Y-axis is the average FSS-7 score and on the X-axis is the perceived effort score for a 25% isometric biceps hold task. Perceived effort score = expected score – given score. Therefore a negative score represents perception of high effort and positive scores represent perception of low effort.

### fMRI

The main effects of hand grip (Bg) were consistent with previous reports using this paradigm [13] and so are not reported in detail here. Regions in which the magnitude of task-related signal increased linearly with increasing hand grip force (Bf) were seen in contralateral M1 (−36, -22, 58), ipsilateral Cerebellum (V) (24, -52, -20), ipsilateral cingulate sulcus (3, 11, 34), ipsilateral primary visual cortex (−6, -94, 4).

### Co-variance of fMRI BOLD measures with Perceived Effort and Central Activation Failure

A strong significant positive correlation was seen between Bf for hand grip and Perceived Effort score indicating that higher the perceived effort greater is the linear increase in activity within pre-SMA (rsq = 0.865) and ipsilateral inferior frontal gyrus (rsq = 0.766) with increasing grip strengths (Table 1, Figure 2). A strong significant positive correlation was found between Bf for hand grip and CAF indicating that higher the central activation failure greater is the linear increase in activity in bilateral striatum (rsq = 0.804, 0.79), ipsilateral dorsal pre-motor cortex (rsq = 0.873) and superior frontal gyrus (rsq=0.913) (Table 1, Figure 3) with increasing grip strengths.

**Table 1:** All regions/clusters are significant at P < 0.05, corrected for multiple comparisons within a priori secondary motor and basal ganglia ROIs (see methods section) (except **\*** P<0.001 uncorrected). All cluster sizes < 50 voxels. The direction of the correlation (+ or -) is given next to the value for the coefficient of determination (r^2^). The values in bold are the cluster/regions that are significant after correction for whole brain multiple comparison.

| Region | Side | Talairach coordinates in MNI space | | | Z-value | Corr ($r^2$) |
| --- | --- | --- | --- | --- | --- | --- |
| | | $x$ | $y$ | $z$ | | |
| **Bf and PE** |  |  |  |  |  |  |
| Pre-SMA | M | 0 | 17 | 46 | 4.38 | (+)0.865 |
| Inferior frontal gyrus* | I | 54 | 41 | -2 | 3.73 | (+)0.766 |
| **Bf and CAF** |  |  |  |  |  |  |
| Superior frontal gyrus | **I** | **21** | **47** | **-2** | **4.85** | (+)0.913 |
| PmD | I | 18 | -22 | 67 | 4.45 | (+)0.873 |
| Caudate (striatum) | **C** | **-18** | **-16** | **19** | **4.00** | (+)0.804 |
| Caudate (striatum) | I | 18 | -25 | 19 | 3.86 | (+)0.79 |

**Figure 2:**
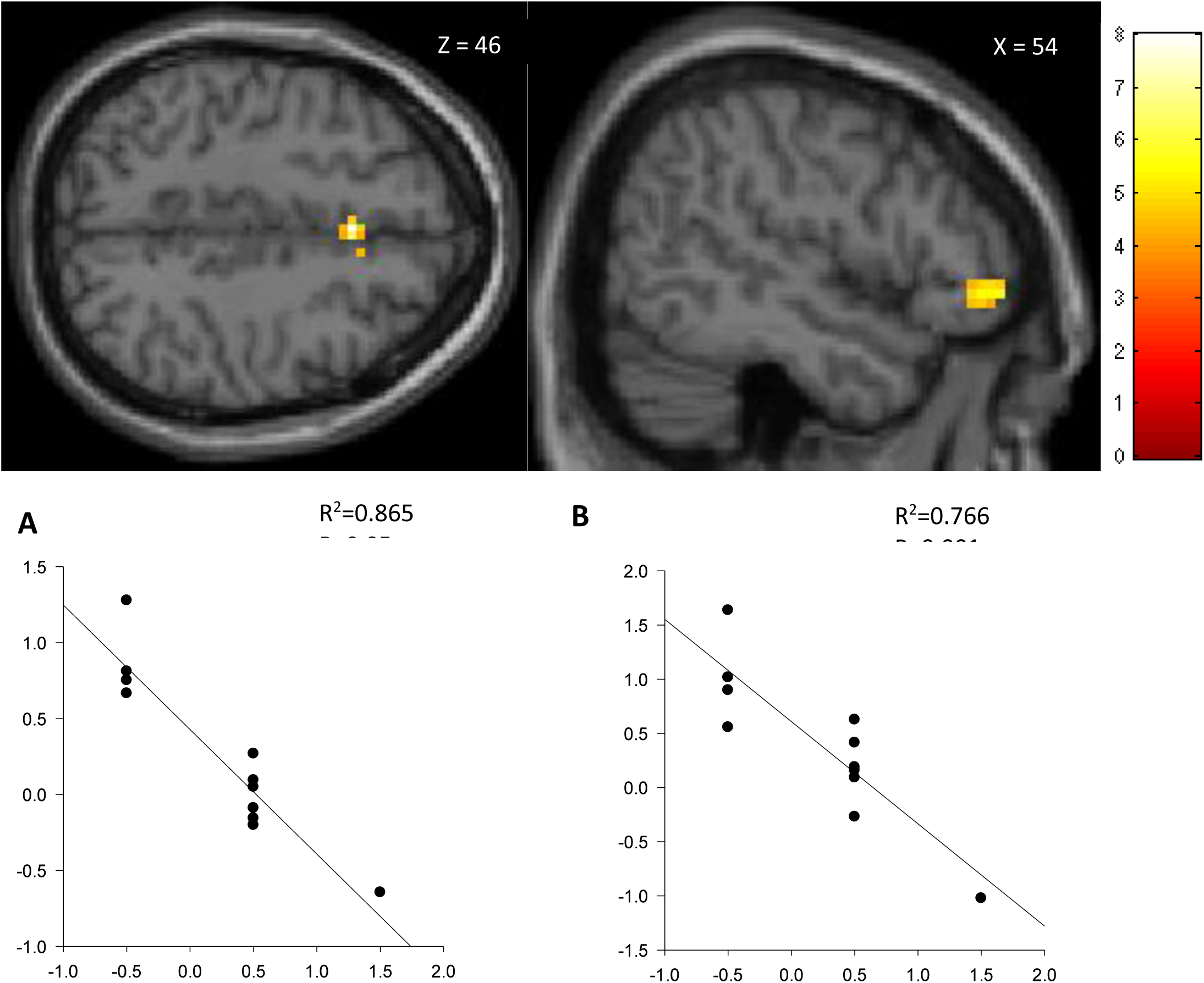
Regions in which increase in BOLD signal with increasing hand grip force (Bf) is linearly correlated with increase in perceived effort is shown overlaid on mean normalised T1-weighted canonical brain. Parameter estimates for the main effect of grip force plotted against Perceived Effort for **A**) pre-SMA (x=0, y=17, z=46) and **B**) ipsilateral inferior frontal gyrus (x=54, y =41, z=-2) are plotted.

**Figure 3:**
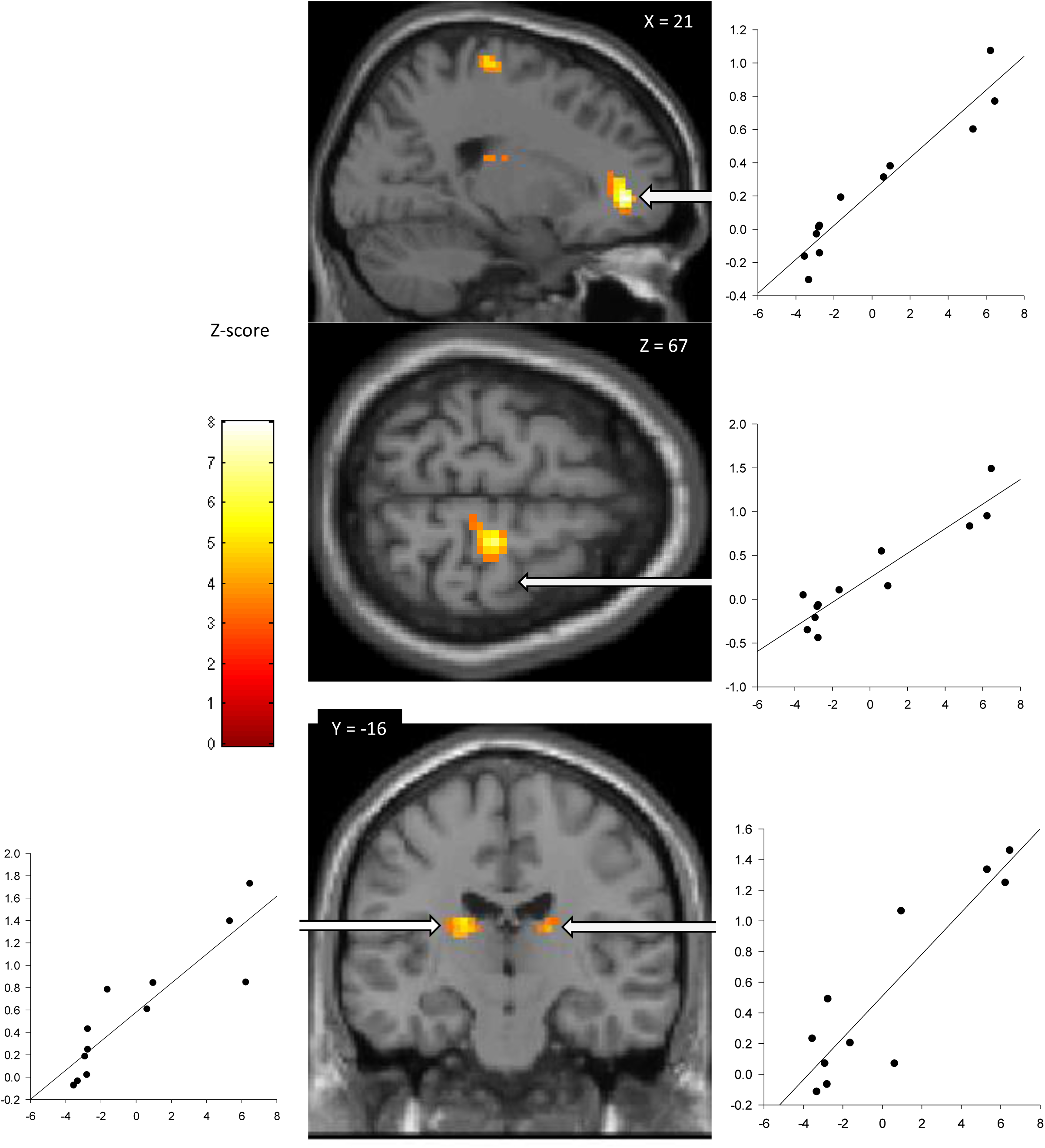
Regions in which increase in BOLD signal with increasing hand grip force (Bf) is linearly correlated with increase in Central Activation Failure is shown overlaid on mean normalised T1-weighted canonical brain. Parameter estimates for the main effect of grip force plotted against Central Activation Failure for **A**) superior frontal gyrus (x=21, y=47, z=-2), **B**) ipsilateral dorsal pre-motor cortex (x=18, y =-22, z=67) **C**) contralateral striatum (x=-18, y=-16, z=-19) and **D**) ipsilateral striatum (x=18, y =-25, z=19) are plotted.

## Discussion

The main findings of this study were 1) high fatigue was associated with high perceived effort 2) higher perceived effort was associated with greater BOLD fMRI activity in pre-SMA and the ipsilateral inferior frontal gyrus with increasing force 3) greater Central Activation Failure was associated with higher increase in BOLD fMRI activity in bilateral caudate, contralateral superior frontal gyrus and pre-motor cortices with increasing force.

Perceived effort, a subjective sense of how much resources one invests in an action, can be classified as a metacognitive act aimed at evaluating one’s own activities. Both SMA and pre-SMA are involved in such metacognition [15,16] and one of the underlying SMA/pre-SMA mediated mechanisms contributing towards assessing one’s own action is sensory attenuation of self-generated force [17]. Sensory attenuation is a unique feature of self-generated forces wherein there is diminished processing of afferent activity arising from muscle contractions that have been voluntarily initiated by the individual. This phenomenon possibly is responsible for the relative effortlessness of performing simple motor actions. In the current cohort of stroke survivors where high perceived effort is seen in high fatigue, greater pre-SMA activity associated with high perceived effort may indicate diminished sensory attenuation i.e. greater sensory processing. Therefore, here we propose a sensory attenuation based mechanism for fatigue in stroke survivors. Increased activation of inferior frontal gyrus (IFG) with higher perceived effort, another finding of this study, may reflect the often seen emotional response that accompanies effort perception. The IFG has been strongly implicated in emotional processing [18] and abnormal activation is seen in many pathological conditions [19].

Central Activation Failure (CAF) is a measure of excitability of the inputs that drive motor cortex output [14]. Motor cortex receives inputs from the basal ganglia [20] and secondary motor areas [21]. The covariance of activity in bilateral striatum and pre-motor cortex with levels of CAF seen in this study further confirms that CAF is a measure of excitability of inputs to the motor cortex. CAF has previously been linked to perceived effort in chronic stroke survivors [9]. Previous studies have also reported encoding of effort levels in the striatum linked to effort-based decision making [22]. The data from this and previous studies suggest a strong link between CAF, perceived effort and fatigue, therefore future studies in stroke survivors would benefit from exploring the direction of causality between fatigue, effort and CAF.

Exaggerated effort perception is a feature of those suffering from central fatigue [23] and has been reported in stroke survivors [6,7]. Quantifying fatigue after stroke is difficult due to its multi-dimensional nature. Questionnaire-based scores [10], the gold standard for fatigue quantification, includes both the sensation of fatigue and the functional impact of fatigue which makes it difficult to interpret. Perceived effort, a variable with fewer dimensions but with strong correlation with fatigue may help better quantify fatigue. In this study we have identified neural correlates of perceived effort and central activation failure in stroke survivors and proposed a possible sensory attenuation based mechanism for effort perception. Understanding the mechanisms of effort perception will help better understand major symptoms like post-stroke fatigue.

